# Comprehensive Proteomic Analysis of HCoV-OC43 Virions and Virus-Modulated Extracellular Vesicles

**DOI:** 10.1101/2024.05.16.594494

**Authors:** Negar Joharinia, Éric Bonneil, Nathalie Grandvaux, Pierre Thibault, Roger Lippé

**Author notes:** Address correspondence to Roger Lippé.

## Abstract

Viruses are obligate parasites that depend on the cellular machinery for their propagation. Several viruses also incorporate cellular proteins that facilitate viral spread. Defining these cellular proteins is critical to decipher viral life cycles and delineate novel therapeutic strategies. While numerous studies have explored the importance of host proteins in coronavirus spread, information about their presence in mature virions is limited. In this study, we developed a protocol to highly enrich mature HCoV-OC43 virions and characterize them by proteomics. Recognizing that cells release extracellular vesicles whose content is modulated by viruses, and given our ability to separate virions from these vesicles, we also analyzed their protein content in both uninfected and infected cells. We uncovered 69 unique cellular proteins associated with virions including 31 high confidence hits. These proteins primarily regulate RNA metabolism, enzymatic activities, vesicular transport, cell adhesion, metabolite interconversion and translation. We further discovered that the virus had a profound impact on exosome composition, incorporating 47 novel cellular proteins (11 high confidence) and excluding 92 others (61 high confidence) in virus-associated extracellular vesicles compared to uninfected cells. Moreover, a dsiRNA screen revealed that 11 of 18 select targets significantly impacted viral yields, including proteins found in virions or extracellular vesicles. Overall, this study provides new and important insights into the incorporation of numerous host proteins into HCoV-OC43 virions, their biological significance and the ability of the virus to modulate extracellular vesicles.

**Importance:** In recent years, coronaviruses have dominated global attention, making it crucial to develop methods to control them and prevent future pandemics. Besides viral proteins, host proteins play a significant role in viral propagation and offer potential therapeutic targets. Targeting host proteins is advantageous because they are less likely to mutate and develop resistance compared to viral proteins, a common issue with many antiviral treatments. In this study, we examined the protein content of the less virulent biosafety level 2 HCoV-OC43 virus as a stand-in for the more virulent SARS-CoV-2. Our findings reveal that several cellular proteins incorporated into the virion regulate viral spread. Additionally, we report that the virus extensively modulates the content of extracellular vesicles, enhancing viral dissemination. This underscores the critical interplay between the virus, host proteins, and extracellular vesicles.

## Introduction

The *Coronaviridae* family is composed of positive strand RNA enveloped viruses that includes four genera, of which the alpha and beta coronaviruses comprise human pathogens. SARS-CoV-2 along with SARS-CoV, MERS and the phylogenetically related HCoV-OC43, belongs to the beta coronavirus genus. HCoV-OC43 is increasingly used as a surrogate for the more virulent SARS-CoV-2 because it is a low-risk and milder endemic respiratory virus, is classified a biosafety level 2 pathogen that makes it easier to handle, and most importantly exhibits a similar dependence on host proteins and sensitivity to pharmacological reagents (1–5). They also share a similar life cycle where the coronavirus spike proteins bind to a cell surface receptor (e.g., ACE2 for SARS-CoV-2 or 9-*O*-acetylated sialic acid for HCoV-OC43), enabling viral fusion with endocytic or plasma membranes depending of the specific coronavirus (6, 7). Gene expression and genome duplication then occur at the endoplasmic reticulum within virus-induced replication compartment (8–11). This is followed by new capsid assembly at the ER-to-Golgi intermediate compartment (ERGIC) and viral egress through the biosynthetic route or possibly the lysosomal pathway. Although these steps of the coronavirus life cycle are known, the molecular machinery driving them remains unclear. As parasitic entities, coronaviruses rely on cellular proteins for propagation, deciphering host-pathogen interactions is a promising approach for targeted therapies (12, 13).

Numerous studies have highlighted changes in the human proteome in response to SARS-CoV-2 and identified host proteins that interact with viral components (14–32). However, it remains unclear which of these proteins are crucial for the virus. Several studies have also used CRISPR-Cas9 to target the human genome and monitor viral outputs (1–3, 33–36). Unfortunately, these large-scale approaches are costly, suffer from limited reproducibility (37, 38) and produce large numbers of candidates making molecular characterization difficult. Our approach, along with others, has focused on analyzing the protein content of viral particles, identifying several host proteins (39–44), some of which are active in viral propagation (45) and have revealed unexpected functions (46, 47). Recent proteomics studies of SARS-CoV-2 virions have suggested that virion-incorporated G3BP1/2 stress granule modulators may play a role in the formation of the viral assembly sites (48). Virion proteomics studies not only provides valuable insights but can also lead to novel broad-spectrum antiviral treatments (42), underscoring the importance of characterizing host proteins in mature virions.

Extracellular vesicles (EVs), including exosomes (30-150 nm) and the larger ectosomes composed of microvesicles and microparticles (50-1000 nm), are secreted by most cells and are involved in various critical functions such as reproduction, development, cancer and diseases (49). EVs also modulate the immune response, play an important role in intercellular communication and are being explored as therapeutic delivery vehicles (49). They contain cellular components such as nucleic acids, proteins, miRNA, lipids and metabolites. There is substantial evidence that EVs are highjacked by viruses or play a role as a cellular defence mechanism (50–55). Altered exosomes have been reported in infections like influenza virus (56), foot-and-mouth disease virus (FMDV) (57), grass carp reovirus (58) and alpha coronavirus porcine epidemic diarrhea virus (PEDV) (59). The modulation of exosomes in COVID-19 patients further supports this concept (60–65). Understanding the interplay between coronavirus and EVs is therefore essential.

This study investigates the protein content of mature HCoV-OC43 extracellular virions and the impact of the virus on EV composition. Our findings reveal that the virus incorporates at least 31 host proteins and significantly alter EVs by promoting the incorporation of 11 high confidence host proteins and the loss of 61 others upon infection. Bioinformatic analyses suggest these changes may alter the functions of EVs. Additionally, a targeted dsiRNA screen validated our proteomics data and indicated that cellular proteins detected in both virions and EVs modulate viral propagation.

## Results

### Preparation of enriched and concentrated virions

We previously reported optimized conditions for propagating and titrating the challenging HCoV-OC43 virus using the fast-growing HRT-18 human cell line (66). We also established serum-free conditions that facilitates virus production to minimize contamination from EVs and serum proteins. This allowed us to obtain sufficient quantity of virus for analysis, but further concentration of virions released in the media was necessary for proteomic studies. Two common methods for this purpose are ultracentrifugation and filtration. Considering the known size range of coronaviruses (80-120 nm) (67), we processed the extracellular media from infected cells by centrifuging at 800 g for 10 minutes at 4° C to remove cell debris, followed by filtration through a 0,45 µm filter to purify the samples further. We then filtered a portion of the cleared supernatants through an Amicon filter with a 100 kDa cutoff, which retains the virus while allowing most proteins to pass. In parallel, we ultracentrifuged another portion of the cleared supernatants at either 20 000, 60 000 or 100 000 *x g* for 60 min and gently resuspended the pellets overnight in 200 µl of PBS. We assessed the infectious viral particles using a sensitive TCID50 immunoperoxidase assay (TCID50-IPA) (66) to determine viral concentration and recovery. According to Table 1, which averages results from three independent experiments, the initial 0.45 µm filtration had no significant effect on viral yield or concentration. In contrast, ultracentrifugation progressively improved viral concentration (up to 16-fold at 100 000 *x g*) with a reasonable recovery (32% of starting infectious particles). However, filtration through the Amicon filter was better with a 29-fold concentration of the virus and a 48% recovery, presumably because this is a milder approach than hard pelleting. Consequently, we continued our research using the Amicon filtration approach to concentrate the virus more efficiently.

**Table 1:** Concentration and recovery of extracellular virus. Supernatants from infected HRT-18 cells were precleared at low speed then filtered (0,45 µm). They were then split and concentrated by two methods, i.e., either through a 100 kDa cut-off Amicon filter or by ultracentrifugation at 20 000, 60 000 or 100 000 *x g*. Viral tit from each step were determined by TCID50-IPA. The initial supernatant was used as the control sample (1x concentration and maximal yield, i.e., 100%). The table shows the average TCID50 and SEM from three independent experiments.

| Method | Sample | Volume | TCID<sub>50</sub>/ml<br>(Mean±SEM) | Concentration<br>(relative to<br>Media) | Total<br>Recovery<br>(%) |
| --- | --- | --- | --- | --- | --- |
| Amicon filter | Media | 3x 60 ml | 3.01 ± 1.65 E+08 | 1 | 100 |
|  | 0,45 µm filtered media | 3x 60 ml | 3.54 ± 1.46 E+08 | 1.2 | 117.5 |
|  | Amicon filter | 3ml | 8.75 ± 2.43 E+09 | 29.0 | 48.2 |
| Ultracentrifugation | Media | 10 ml | 8,75 ± 2.43 E+07 | 1 | 100,0 |
|  | 0,45 µm filtered media | 10 ml | 8,75 ± 2.43 E+07 | 1,0 | 100,0 |
|  | Pellet 20K x g | 0.2 ml | 3.01 ± 1.65 E+08 | 3.4 | 6.9 |
|  | Pellet 60K x g | 0.2 ml | 5.19 ± 1.13 E+08 | 5.9 | 11.9 |
|  | Pellet 100K x g | 0.2 ml | 1.40 ± 0.77 E+09 | 16.0 | 32.0 |

While our samples were devoid of contaminating serum, both uninfected and infected cells can release vesicles in their extracellular environment. It was therefore critical to separate virions from EVs, particularly for the similarly sized exosomes. To first assess the presence of EVs, we monitored the presence of the CD9, CD63 and CD81 classical exosome markers by Western blotting (49, 68, 69). Using total lysates as positive controls, we found evidence that HRT-18 cells express CD9 and CD63 (fig. 1A, total cell lysates). Under these conditions, we also detected CD9 in the media (fig. 1A, media). This remained true when the media was concentrated at 100,000 *x g* (fig. 1A, pellet), the prototypic condition to bring down EVs (69). Note that the anti-human CD81 antibody was functional as it detected the protein in the unrelated Vero cell line (fig. 1B). Interestingly, the virus had no noticeable impact on the expression level or secretion of the exosome markers.

**Figure 1:**
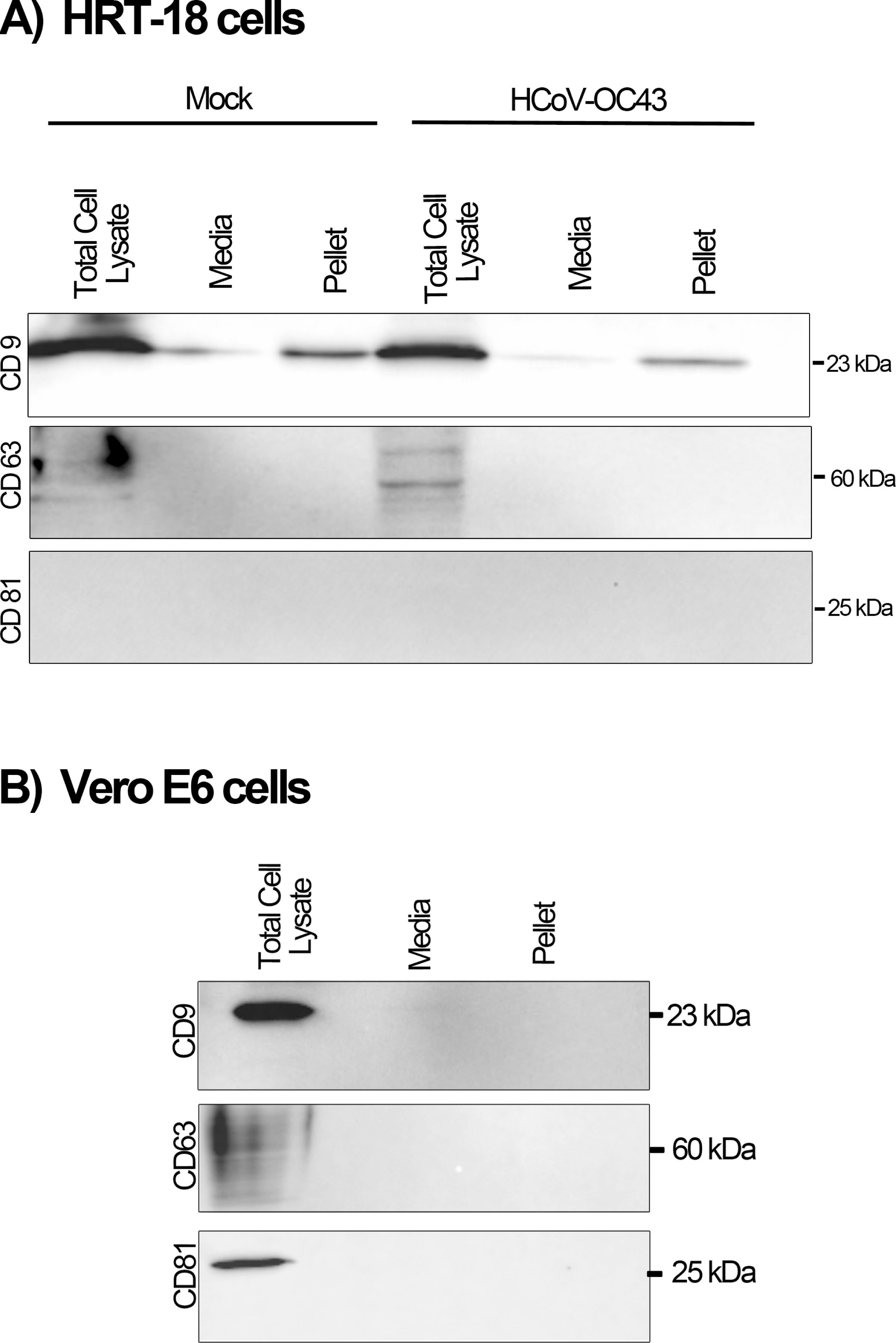
A) Probing of exosome makers in HRT-18 cells. HRT-18 cells were infected with the HCoV-OC43 VR-1558 virus at a MOI of 0.1. Infected cells were incubated in SFM-OptiPRO to avoid potential contamination by exogenous exosomes. Three days post infection, supernatants were collected, filtered with a 0,45 µm filter and ultracentrifuged at 100 000 x g for 1h. Exosome markers CD9, CD63, and CD81 were detected by Western blot in total cell lysates, the 0,45 µm filtered media as well as the ultracentrifuged pellets on both mock and HCoV-OC43-infected samples. Molecular weights are indicated to the right of the blots. **B) Validation of the CD81 antibodies.** Vero E6 infected cells and corresponding supernatant and high-speed pellets were used as control for the CD81 antibodies. All blots are representatives of three independent experiments.

Separating EVs from virions has been a critical issue. Recently, Dogrammatzis and colleagues reported that an iodixanol/sucrose density gradient efficiently separates HSV-1 virions from EVs (70). We thus probed whether such an approach could also resolve HCoV-OC43 virions and EVs. HRT-18 cells were thus infected at a multiplicity of infection (MOI) of 0.1 in serum-free conditions and the media harvested three days post-infection. The samples were next concentrated on Amicon filters, which retains the virions but also the similarly sized or even bigger EVs, and loaded them onto 8-25% iodixanol/sucrose gradients. After centrifugation, eighteen fractions were collected from top to bottom and monitored by Western blotting by probing the N viral protein or the CD9 and CD63 exosome markers expressed in HRT-18 cells. As positive control, total cell lysates were monitored while an equal number of non-infected cells was used as negative control. Figure 2A shows that although some HCoV-OC43 N protein was trailing across the gradient, it principally accumulated in the denser fractions (fractions 17-18). As expected, no evidence of the N viral protein was found in the mock samples. Most interestingly, CD9 and CD63 peaked in the middle of the gradient (fractions 4-11) and were barely detectable in the virion fractions. As noted above, the expression level and density of the exosome markers appeared unaltered by the virus (fig. 2A; compare mock and infected samples). Based on these criteria, we deemed it possible to efficiently separate HcoV-OC43 virions from the EVs that may be present in the media.

**Figure 2:**
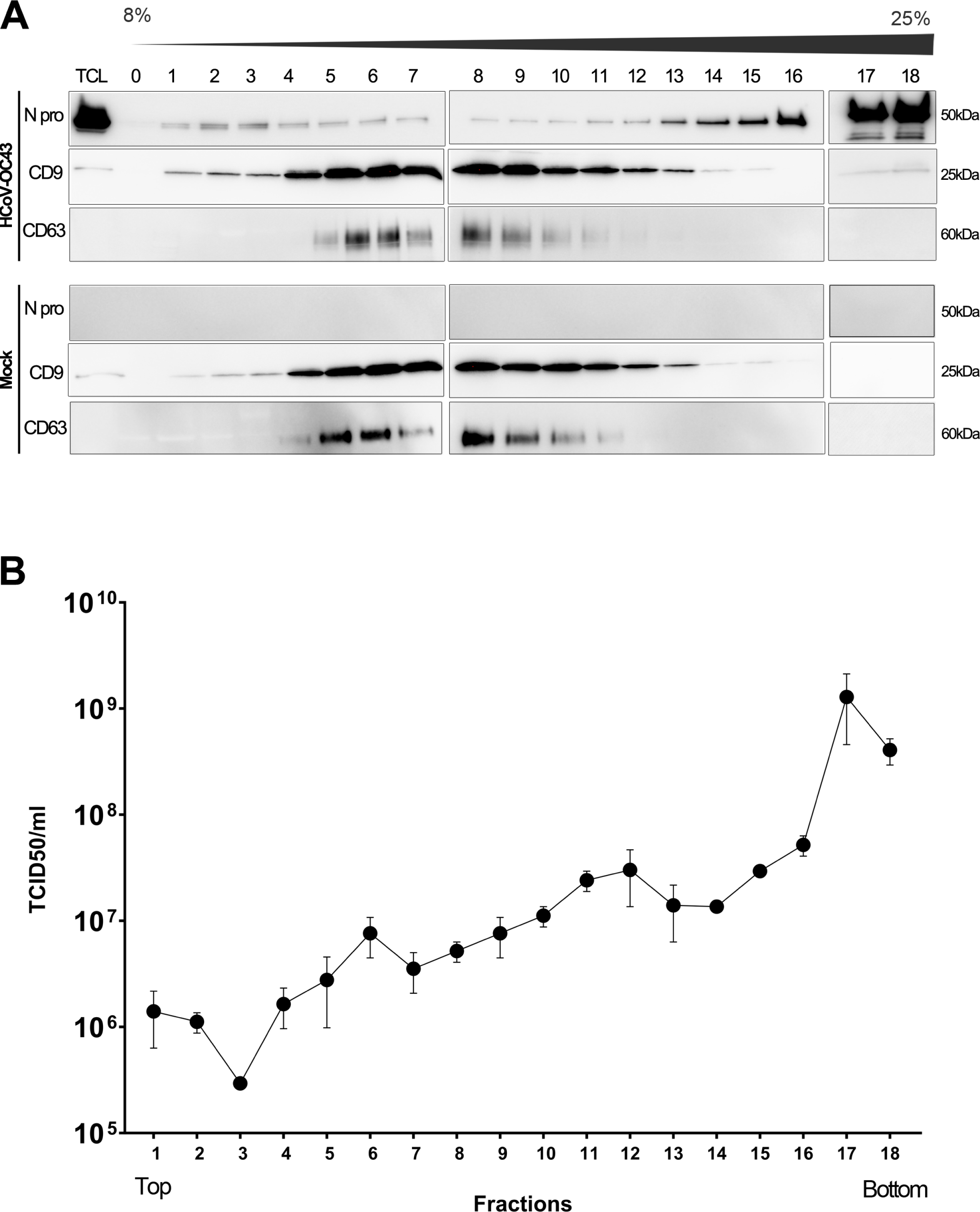
Separation of EVs and virus on density gradients. **A)** HRT-18 supernatants from mock-infected or HCoV-OC43-infected cells incubated in SFM OptiPRO media were collected three days post-infection, 0,45 µm filtered then passed in 100 kDa Amicon Millipore filters. The concentrated samples were then centrifuged on continuous 8 to 25% iodixanol/sucrose density gradients and 18 fractions were collected from the top (fraction 1) to the bottom (fraction 18). For each fraction, 50 µl were resolved on 5-20% SDS-PAGE gradient gels. Total cell lysates were used as positive controls. The fractions were probed for exosome markers (CD9, CD63) and HCoC-OC43 (N protein). The blots are representative of three independent experiments. **B)** Fractions from the above density gradients were tittered by TCID50-IPA. Error bars present SEM from the three independent experiments.

Given our interest to characterize the protein content of mature and functional virions, it was critical to assess the infectivity of the collected fractions. We thus monitored the same fractions by TCID50-IPA using a monoclonal antibody against the surface spike protein (66). The data indicate that infectious particles followed a similar pattern as the N-protein and were far more abundant in fractions 17 (fig. 2B). As fraction 18 was the bottom of the density gradient (i.e., it included the pellet), this suggested it may be difficult to fully resuspend the virus, in agreement with the reduced recovery of the virus by ultracentrifugation.

Given the efficient separation of HcoV-OC43 virions and EVs, we arbitrarily pursued our analyses using fractions 6 (EVs) and 17 (virions). To further validate their identity and purity, we first analyzed them by silver staining by comparing samples from mock-treated or infected cultures. As anticipated, several bands that putatively corresponded to viral proteins, based on their molecular weights, were readily detected in fraction 17 from HcoV-OC43-infected samples (fig. 3). In contrast, the results point to the near absence of proteins in fraction 17 from mock-infected cells, hinting that it lacked virions but also EVs or other cellular components. Meanwhile, several proteins were shared between the exosome fractions, with some discernible differences between the mock and infected samples. Overall, this further confirmed the proper separation of virions from EVs and the presence of virions in fraction 17. To orthogonally validate these findings, we next examined fractions 6 and 17 by negative staining and electron microscopy. For the uninfected sample, small smooth vesicles with a darker center were present in fraction 6 (fig. 4), as expected for exosomes (71). Upon infection, these vesicles exhibited a different aspect with the loss of the darker center, suggesting that the virus somehow altered them. For fraction 17, typical spiked structured were present in the infected samples but absent in the mock sample. Importantly, all the fractions were relatively free of contaminants. Size measurements of 15-20 particles per sample indicated that the samples in fraction 17 contained virion-size particles (78.7 nm + 1.5 (SEM)) and that fraction 6 fitted the expected size of exosomes (33.2 nm +1.0 (SEM)) with no evidence of larger ectosomes. Moreover, virions were statistically larger than exosomes (p<0.05). We thus concluded that both virions and EVs were relatively devoid of contaminants and minimally, if at all, cross-contaminated with one another.

**Figure 3:**
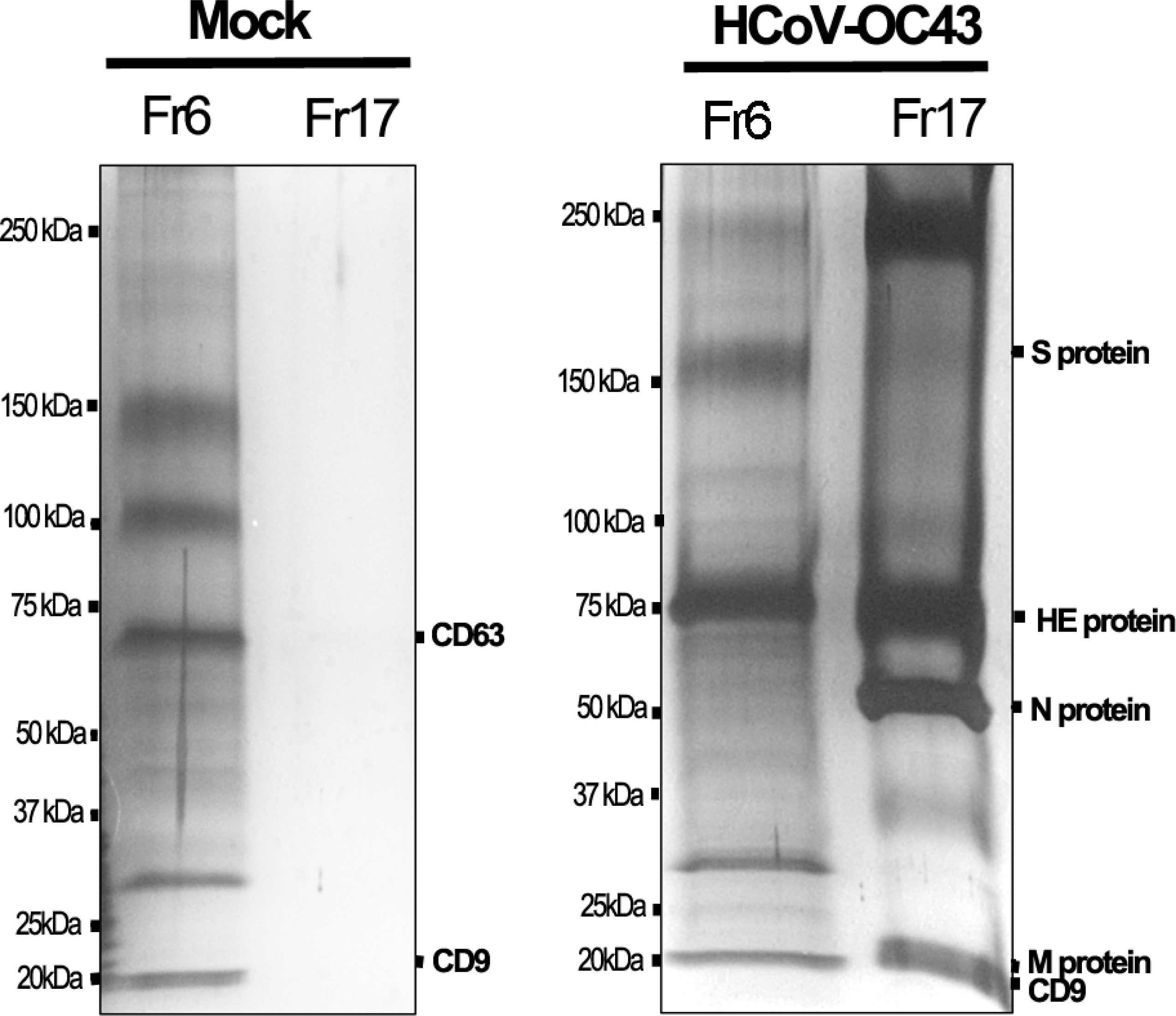
Distinct silver staining pattern of mock and HCoV-OC43-infected fractions. EVs and virion-containing fractions collected from mock and HCoV-OC43 infected samples were loaded on 5-20% SDS-PAGE gradient gels and silver stained. The expected migration of the HCoV-OC43 S, N, M, HE proteins and predicted exosome CD9 and CD63 markers are indicated. The data presented here is representative of 3 independent experiments.

**Figure 4:**
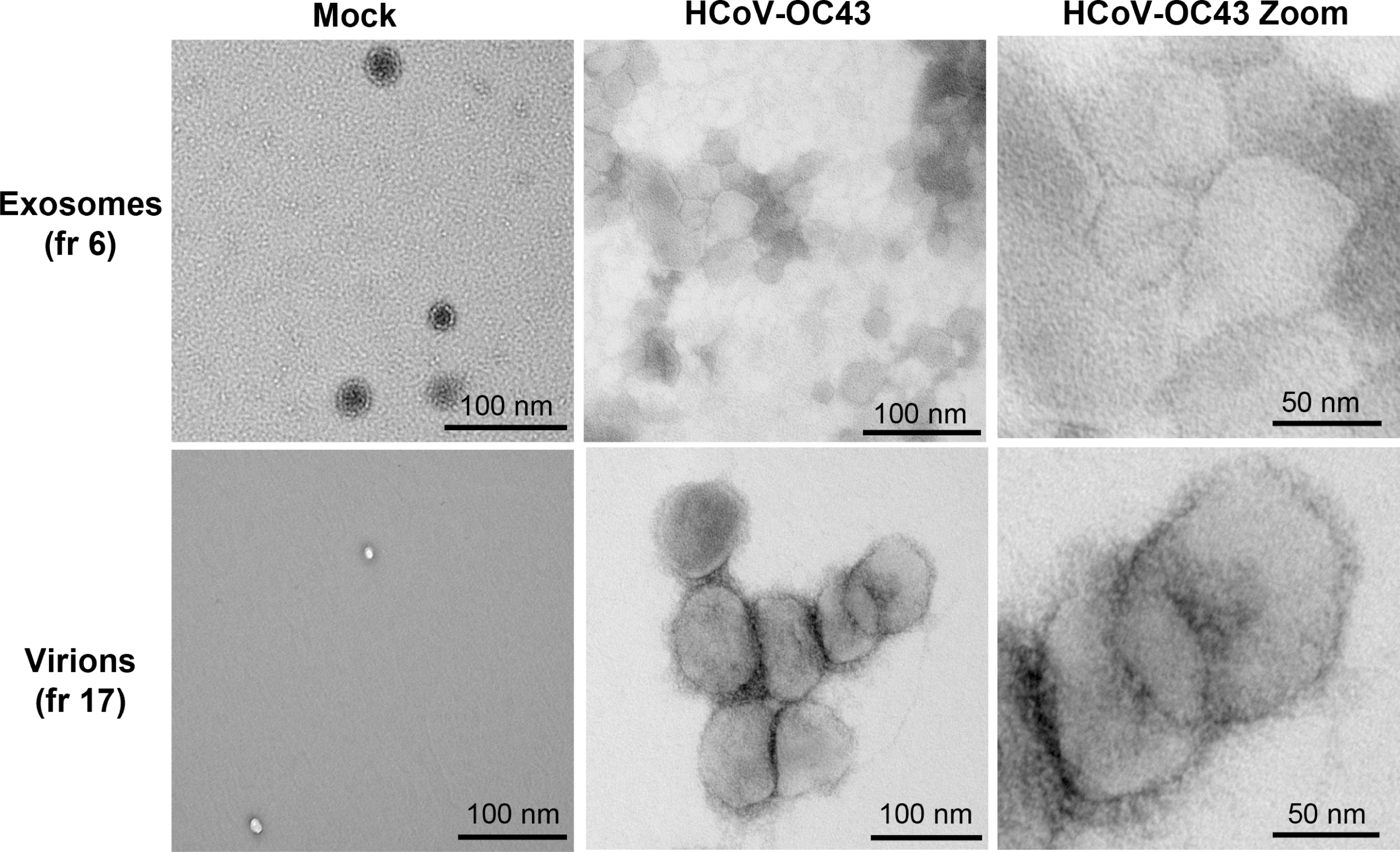
Transmission election microscopy imaging of exosome and viral fractions. Exosome (fractions 6) and virion fractions (fractions 17) from mock or infected cells were inactivated with 0.8% PFA then examined by negative staining TEM The data presented here is representative of 3 independent experiments.

### Proteomics of virions and EVs

In view of the above results, we analyzed by mass spectrometry (MS) fractions 6 and 17 from three independent experiments derived from either mock or infected cells. Using a combined Homo sapiens and a HcoV-OC43 protein database, between 25 894 and 32 379 total spectra were detected for each sample (fig. 5). Using stringent conditions (95% protein & peptide thresholds; 2 peptides minimum per protein), 414 different proteins were found in the virion fractions, as well as 613 and 617 proteins in the EVs from mock and infected fractions, respectively. Meanwhile, 231 proteins were identified in fractions 17 from uninfected cells. To limit putative false positives, only the hits that were reproducibly detected in the triplicates were considered (bold blue numbers in fig. 5). This led to 385 and 335 total proteins (viral and cellular) in mock and infected EVs, respectively (fractions 6, mock/infected) (fig. 5). We also found 163 distinct proteins associated with virions (fraction 17, infected) and 61 in the negative control (fraction 17, mock). Interestingly, four of the five known HcoV-OC43 structural proteins were found in the virions (fraction 17, infected cells), including the spike (S), nucleoprotein (N), membrane (M) and hemagglutinin esterase (HE). In contrast, the envelope (E) viral protein eluded MS detection. As noted above, traces of viral proteins were detected in the EVs isolated from infected cells, but these were strongly enriched in virions (Table 2). The HCoV-OC43 nonstructural protein 2A was present in virions but not EVs, the significance of which is not clear.

**Figure 5:**
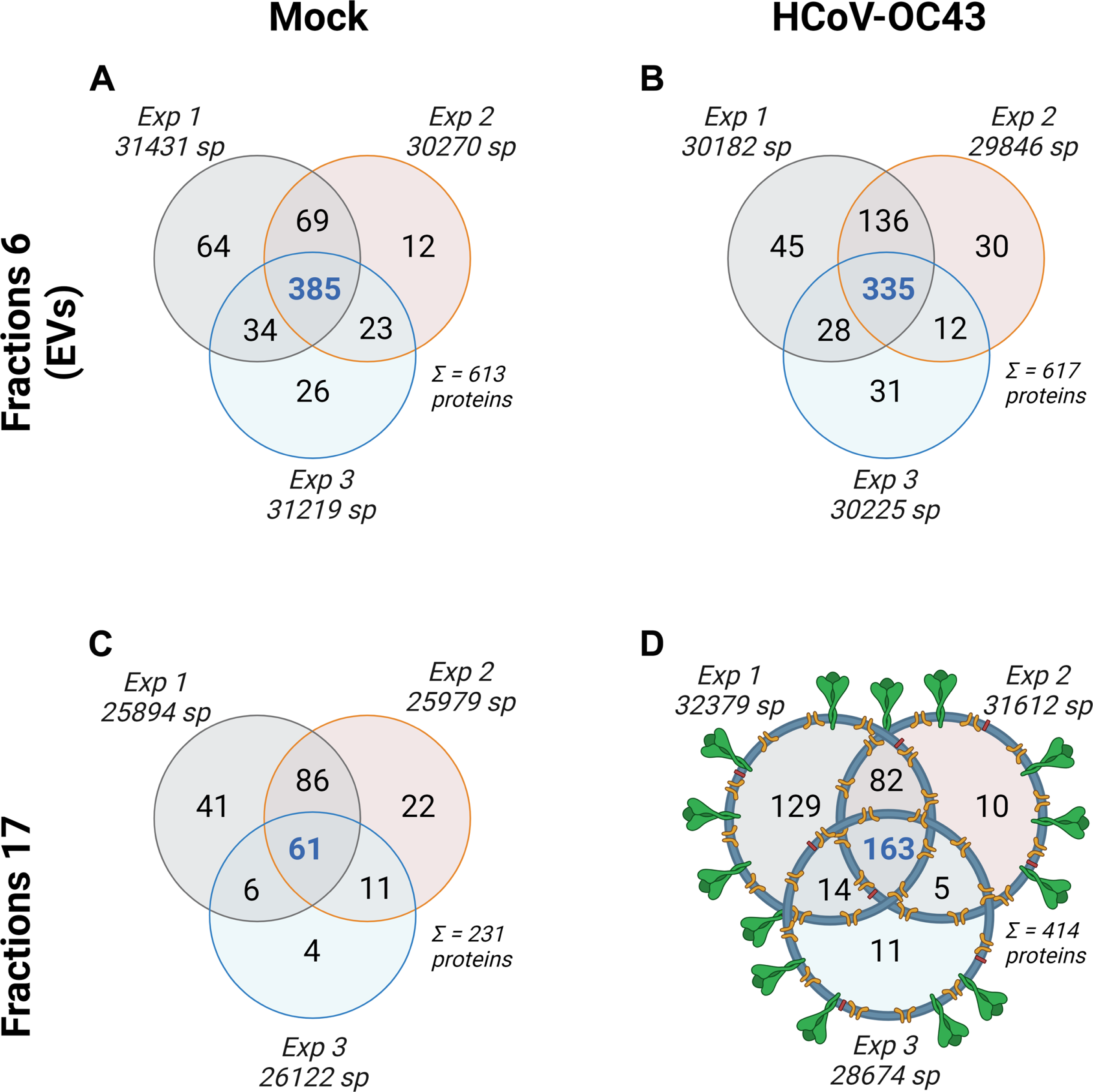
Total protein content of virions and exosome fractions. The figure represents all the proteins (viral and cellular) identified in fractions 6 (exosomes) isolated from either **A)** mock or **B)** infected-cell derived supernatant, as well as fractions 17 from **C)** control uninfected cells derived supernatant and **D)** virions. The total numbers of proteins identified and spectra (sp) are indicated. Proteins that were found in all three replicates are indicated in bold blue. This figure was generated with Biorender.

**Table 2:**
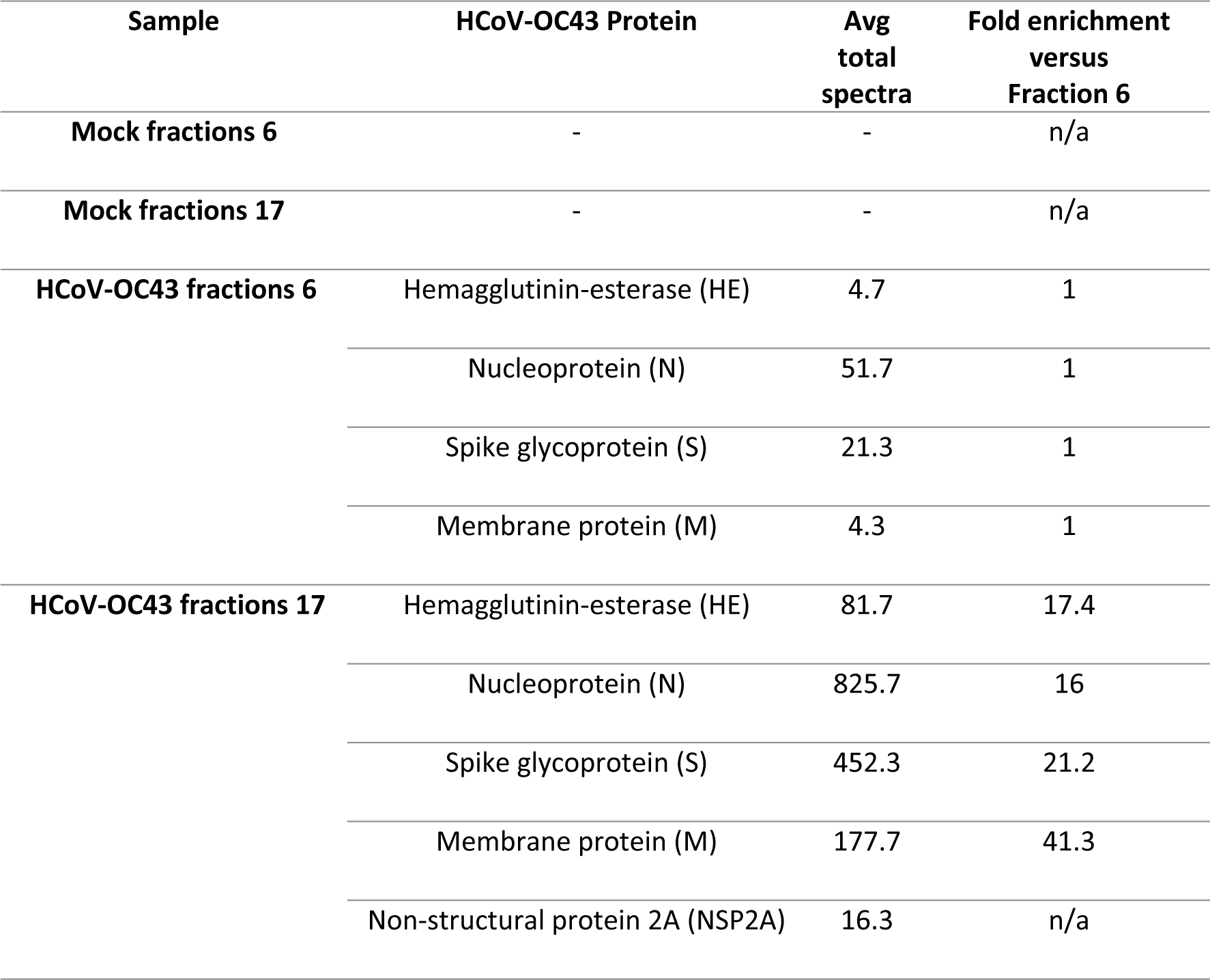
Viral protein enrichment in diverse fractions. The presence of viral proteins in the different samples is shown (average total spectra from three independent experiments). Note the absence of viral proteins in the non-infected samples. An enrichment of viral structural proteins (ratio fractions 17/fractions 6) is evident when compared to the infected exosomes. n/a: not applicable.

### Data Curation

Before proceeding to functional assays, we deemed it essential to validate our MS results. In this study, the protein content of the EVs and viral fractions were derived from Scaffold, a commercial proteomics tool. A limitation of this software is the inclusion of hits in Venn diagrams when a single sample among the triplicate meets the two-peptide minimal criteria (Fig. 5). We therefore hand curated our findings to limit ourselves to the samples that were reproducibly identified and met our stringent criteria in each triplicate. While this potentially eliminated proteins of interest, it enhanced data quality by removing putative false positives. Most importantly, this step had no impact on the viral proteins listed above but reduced the cellular content of the samples to 261 (mock EVs), 211 (infected EVs), 30 (control mock fraction 17) and 87 (virions) (fig. 6A, B; Supplementary Tables). A single protein (ARF4) was common among the unique proteins found in infected EVs and those in virions, highlighting the purity of the samples (fig. 6C).

**Figure 6:**
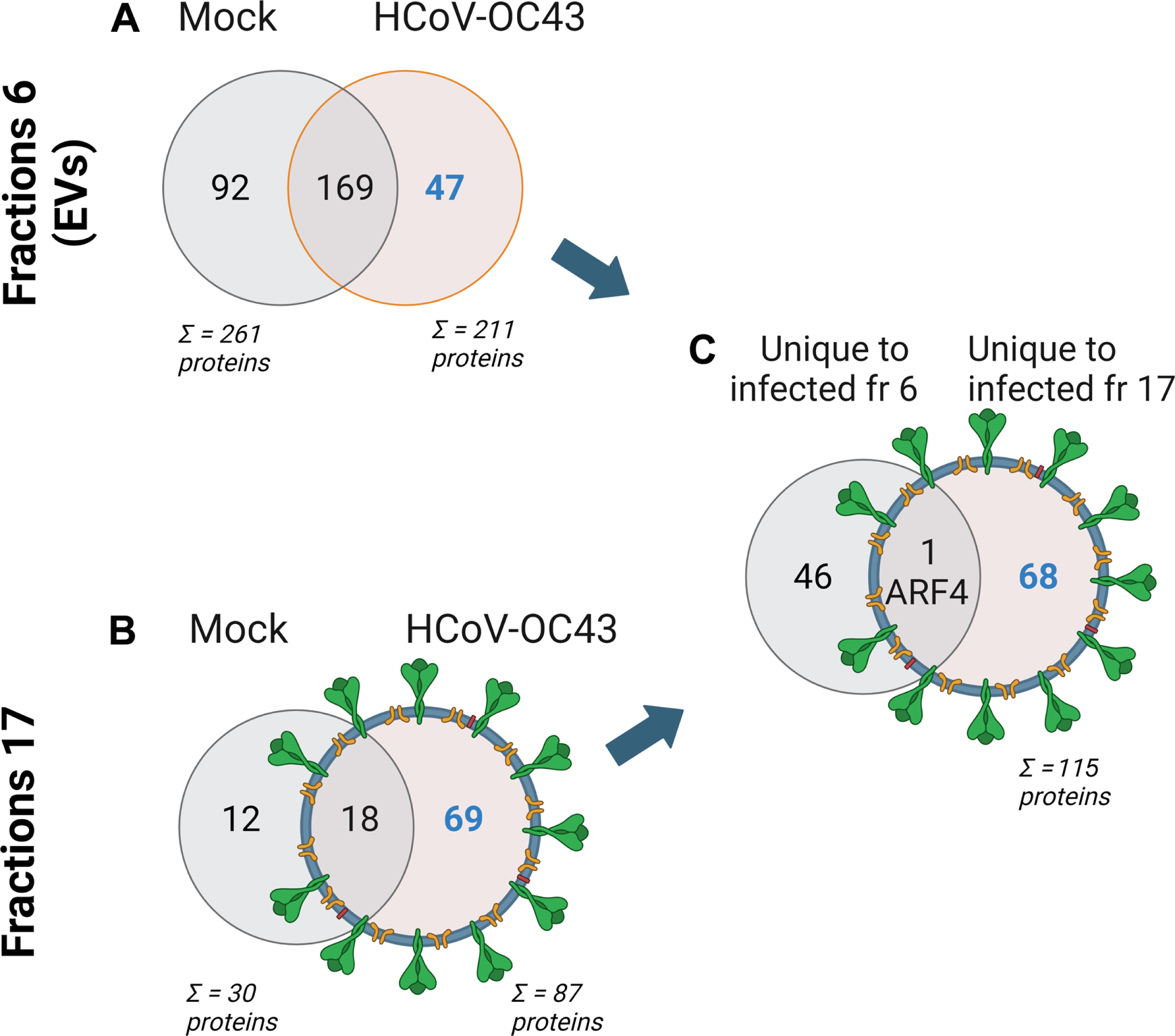
Curated host proteins. **A, B)** Host proteins that were reproducibly found in the three independent experiments were hand curated to exclude samples that did not reach the minimum two peptide threshold in all the triplicates. **C)** Proteins that were either unique to infected exosomes or virions were compared. This figure was generated with Biorender.

As an additional step to ensure the stringency of our protein hits, we considered the presence of contaminants that may co-sediment with the EVs and virions. We therefore ran our hand-curated hits against the Contaminant Repository for Affinity Purification (CRAPome) database (72). This left us with 31 high confidence virion-associated cellular proteins that were absent in the corresponding fraction of the uninfected control samples, 11 proteins unique to EVs derived from infected cells and 61 host proteins unique to mock EVs (fig. 7A; those high confidence hits are shown in bold in the Supplementary Tables). In addition, 93 high confidence cellular proteins were common to both types of EVs. Among them, 82 proteins were detected at similar levels in the mock or infected EVs (fewer than 2-fold differences in average total peptides; fig. 7B). In contrast, nine proteins (ALPP, DIP2B, HSPG2, OLFML3, PLOD3, PLXNB2, SDCBP2, TINAGL1, TSPAN14) were enriched in the mock EVs and two (INS, TPP2) were more abundant in the infected EVs (fig. 7B). Importantly, we detected CD9 and CD63 but also CD81 in EVs from mock infected cells (mock fraction 6; fig. 7A). Interestingly, CD63 was only present in the EVs from the uninfected cells, as agreement with our silver staining results (fig. 3). The EVs were also positive for the TSG101, VPS4A and annexins A2 & A11 EV markers and lacked common contaminants such as APOA, as recommended by the MISEV consortium (69). Aside from annexin A11, none of these EV markers were found in the virion fractions. This once again confirmed the minimal presence of EVs in the viral fractions. Altogether, these results implied that the virus incorporated several cellular proteins, promoted the incorporation of 11 proteins in EVs and excluded 61 other proteins normally present in EVs produced by non-infected cells. This strongly indicated that the virus impacted the content of EVs, perhaps justifying the change of morphology seen above by electron microscopy.

**Figure 7:**
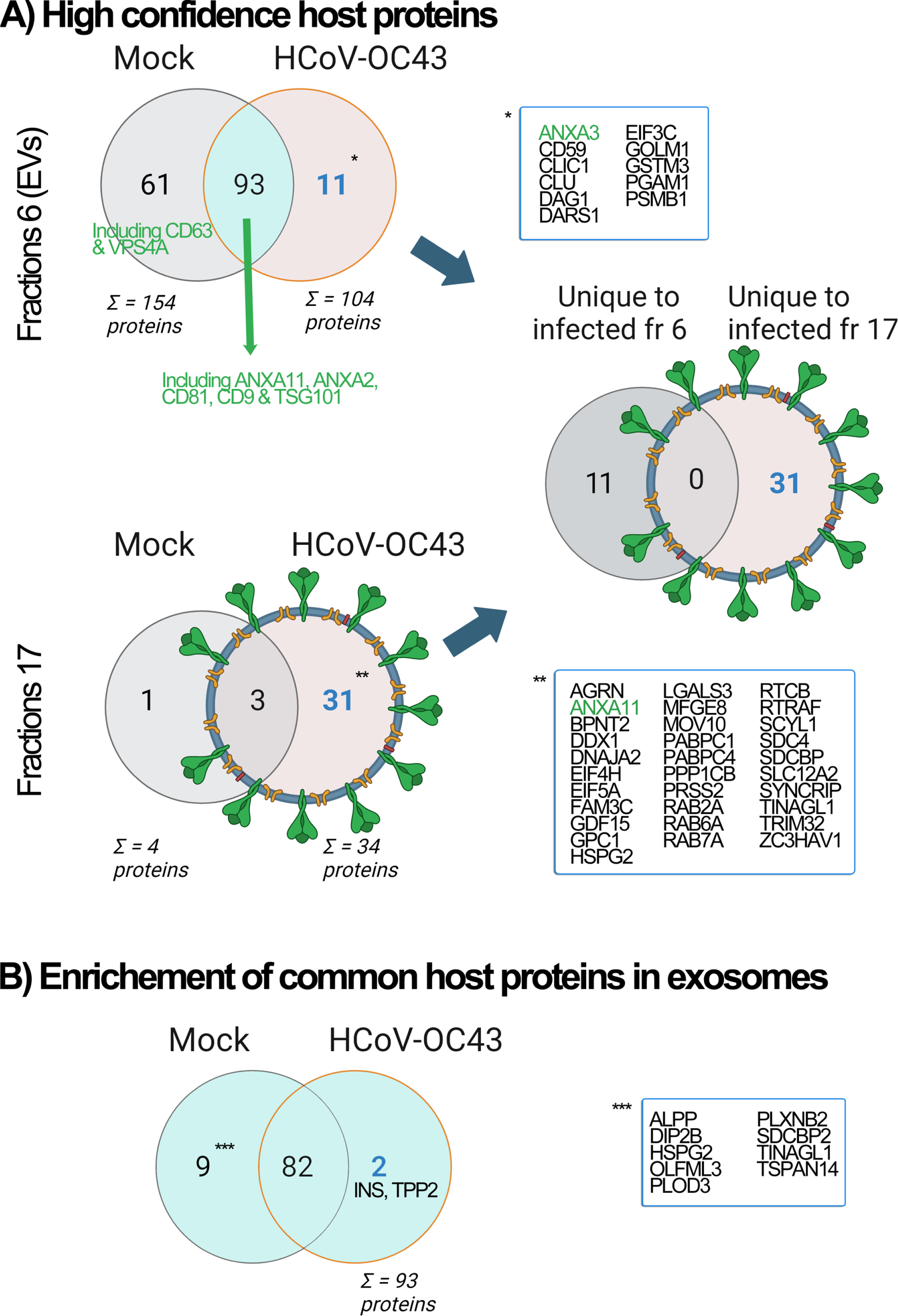
High confidence host proteins. **A)** The Venn diagrams show the hand curated host proteins that were reproducibly found in the three independent experiments but were absent in the CRAPome database (potential contaminants). **B)** The 93 proteins that were common to mock and infected EVs were analyzed for their potential enrichment in either sample. This figure was in part generated with Biorender. EV markers are shown in green.

### Functional Analyses

To decipher the potential role of the high confidence hits identified in this study, we focused on the host proteins incorporated in the virions and EVs and analyzed them with the Protein Analysis Through Evolutionary Relationships (PANTHER) software, which regroups proteins according to their common functions (73, 74). This revealed the presence of virion-associated host proteins involved in RNA metabolism (DDX1, MOV10, PABPC1, PABPC4, SYNCRIP), protein modifying enzymes (PRSS2, PPPACB, SCYL1, TINAGL1), vesicular transport (RAB2A, RAB6A, RAB7A), cell adhesion (AGRN, HSPG2), metabolite interconversion (BPNT2, MFGE8), translation (EIF4H, EIF5A) and nine other functional groups (fig. 8A). Meanwhile, HCoV-OC43 impacted the total protein composition of the EVs with a bias towards protein modifying enzymes, cell adhesion and metabolite interconversion at the expense of protein-binding modulators, cytoskeletal components, extracellular matrix and chromatin-binding. This impact was evident when specifically looking at the cellular proteins that were unique to infected cell-derived EVs (fig. 8B, C), including novel proteins modulating metabolic interconversion (GSTM3, PGAM1), translation (CLU, EIF3C, DARS1), calcium-binding (ANXA11), cell adhesion (DAG1), protein modification (PSMB1) and transporters (CLIC1).

**Figure 8:**
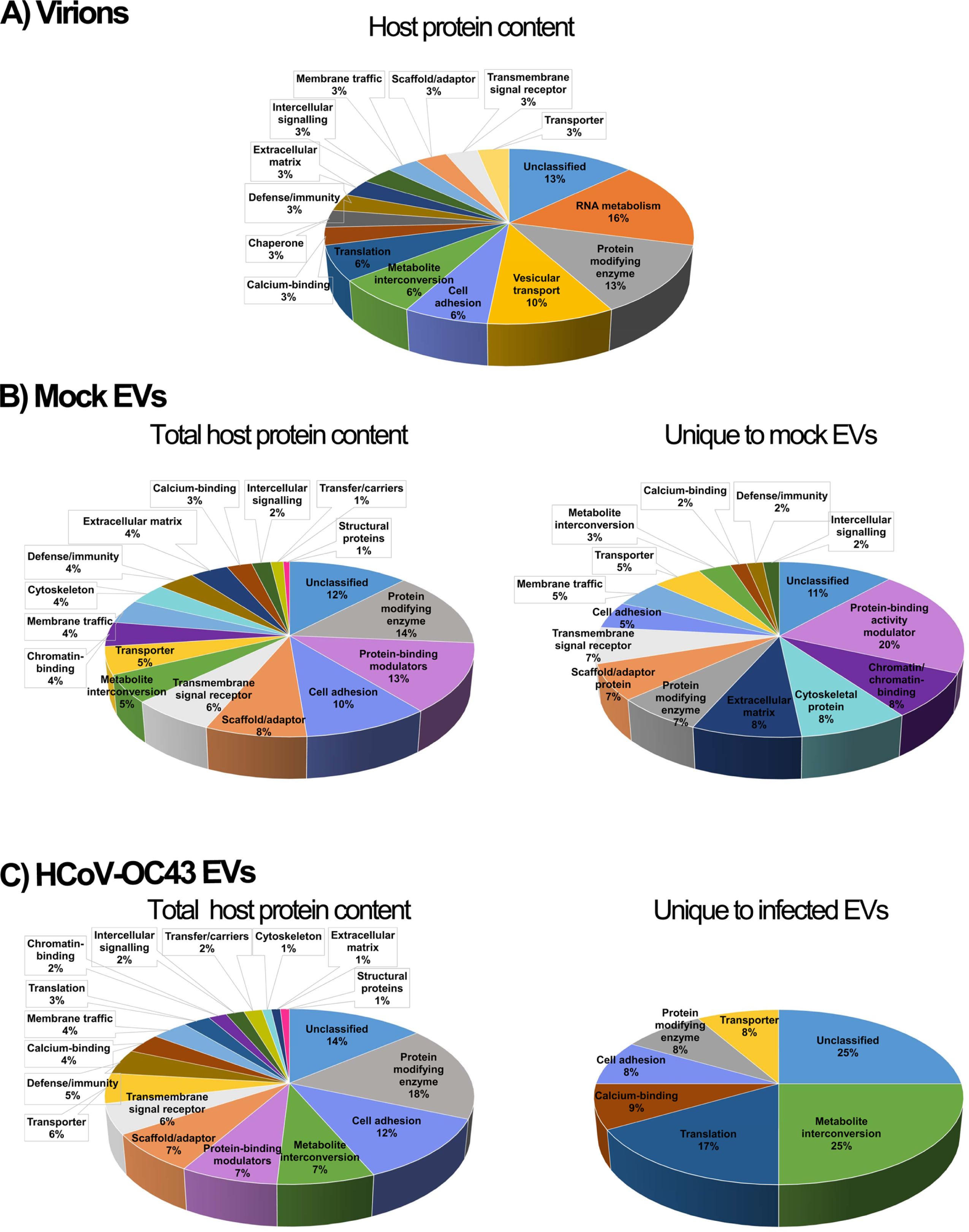
PANTHER analysis of the high confidence host proteins. The role of the cellular proteins associated with the **A)** virus, **B)** mock EVs and **C)** infected EVs were probed using the PANTHER software. The pie charts include 31 virus-specific cellular proteins, 154 host proteins for uninfected exosomes (of which 61 are unique to that sample) and 104 host proteins for infected exosomes (11 unique proteins to that sample). Only reproducibly detected, hand-curated and CRAPome negative proteins were considered.

To validate our data, we performed a select dsiRNA screen to evaluate the ability of proteins identified with PANTHER to modulate HCoV-OC43 propagation. Specific targets were picked through the Ingenuity Pathway Analysis software (Qiagen) based on their interconnectivity with other components, estimating that if these central proteins play a role, the neighbouring components in the same pathways may also be involved. Hence, the HRT-18 cells were pretreated for 24 h with dsiRNA targeting 18 proteins (14 targets associated with the virions and 4 specific to infected EVs) or a control dsiRNA (NC1) then infected with HCoV-OC43. A further 48 h later, cell-associated viral yields were measured by TCID50-IPA. Figure 9A shows that 11 of the 18 tested dsiRNAs (61% positive hits) statistically reduced viral titers, with four dsiRNAs reducing them by at least 75%. This included virions-associated proteins (RAB2A, SCYL1, PABPC4, PPP1CB, PRSS2, SYNCRIP, DDX1, RAB7A and MFGE8) or EV proteins from infected cells (EIF3C, CLU). To ensure these dsiRNAs did not indirectly dampen viral propagation by impacting the integrity of the cells, cell viability was monitored. Fortunately, incubation of the cells with dsiRNA for 72 h did not have any major impact on cell viability, with the exception of EIF3C that reached statistical significance with 20% mortality and oddly TINAGL1 with a mere 10% mortality. However, these rates seemed negligible compared to the 91.8% reduction in viral yields for EIF3C while the 17.9% reduction for TINAGL1 mirrors the reduction in cell viability (fig. 9B). This indicated that many of the high confidence hits identified by MS indeed regulated the HCoV-OC43 life cycle.

**Figure 9:**
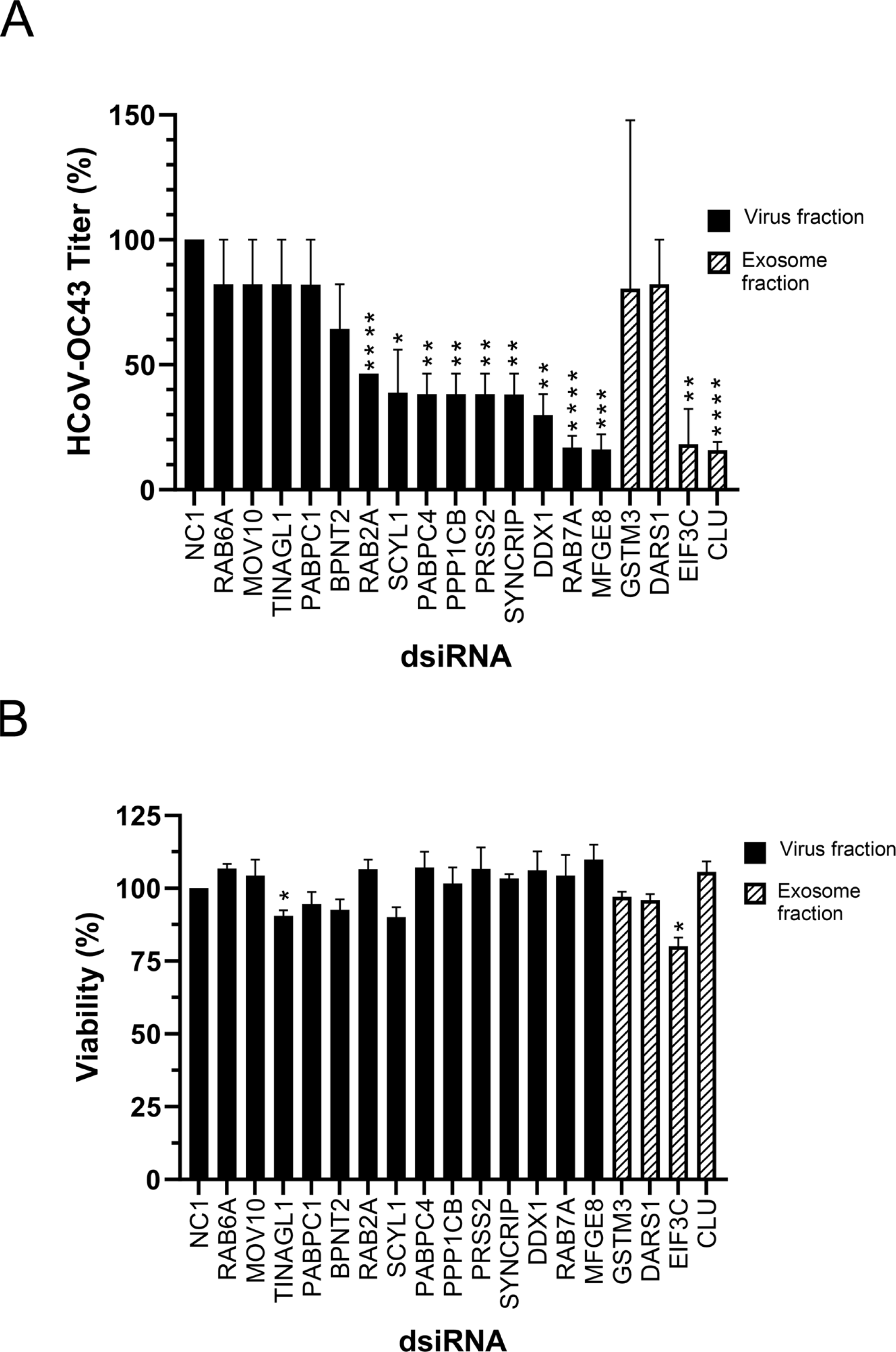
Impact of select protein hits on HCoV-OC43 propagation. **A)** HRT-18 cells were transfected with 100 nM dsiRNAs for 24 h, then infected with HCoV-OC43 at an MOI 0.1 for 48 h. Thereafter, cell-associated virions were collected and HCoV-OC43 quantified using the TCID50-IPA method. **B)** Cell viability was determined using 10% alamarblue, 72h after transfection with 100 nM dsiRNAs. In both panels, the black bars refers to the host proteins found in the virion fractions while striped bars are the host proteins found in the exosome fractions. The error bars represent the means and SEM from three independent experiments. T-test was to analyze the significance of the data (*: p < 0.05; **: p < 0.01; ***: p<0.001, ****: p< 0.0001). All dsiRNAs were normalized to the NC1 control.

## Discussion

Deciphering the implication of cellular proteins in the propagation of human pathogens offers a promising strategy to identify new therapeutic targets. For coronaviruses, particularly SARS-COV-2, numerous studies have utilized genome-wide CRISPR screens, explored the viral interactome and analyzed proteomic changes in infected cells (1–3, 14–36). In contrast, our lab and others have developed a targeted proteomic pipeline that scrutinizes the protein content of highly purified viral particles (39–44). Although this approach is less explored, it is pertinent as many viruses incorporate host molecules, some of which are biologically active (45–47, 75–79). Our focused proteomic method has several advantages, including a limited number of identified targets that simplifies subsequent functional analyses. This approach typically yields a high rate of positive hits that influence virus behaviour (61% in this study), which is significantly higher than results from broader genomic screens. An important limitation, however, is the requirement for high purity samples prior to MS analysis, which can be compromised by abundant serum proteins and EVs in the growth media. Our protocol addresses this by using serum-free conditions to cultivate the infected cells (66), thereby reducing unwanted proteins and EV contamination. The challenge of distinguishing viral particles from EVs, which share similar biochemical properties, is significant given both entities’ interaction with each other and their composition are crucial areas of interest. Kalamvoki and colleagues provided an innovative solution for separating virions from EVs using density gradient separation (70). By integrating their method with ours, we achieved high purity samples suitable to MS analysis. The purity of these samples was confirmed through various techniques, including silver staining, Western blotting, electron microscopy and the detection of specific protein markers. For instance, our MS data confirmed the presence of typical markers (CD9, CD63, CD81, TSG101, VPS4A and annexins A2 & A11) in the EV fraction and the absence of common contaminants such as APOA, as recommended by the MISEV consortium (69). Aside from annexin A11, none of these EV markers were found in the virion fractions. Furthermore, the EV and virion protein content did not overlap, with only a sole common protein (ARF4) among the 115 different host proteins initially identified and no shared proteins once “putative contaminants” were removed via the CRAPome database. Finally, while trace amounts of virus were found in the EV fractions by Western blotting and MS, these were strongly enriched in the viral samples. We are thus confident that the samples are of significant and proper purity.

The present approach revealed the putative presence of hundreds of cellular proteins in virions. By focusing on the hits that were reproducibly found in the three independent experiments and removing potential contaminants through CRAPome, a short list of 31 high confidence host proteins were identified. These proteins primarily modulate RNA metabolism, protein modifying enzymes, vesicular transport, cell adhesion, metabolite interconversion and translation. It should be noted that the high stringency applied to our analyses likely excludes some positive hits that should not formally be ruled out. HCoV-OC43 also caused the exclusion of 61 host proteins in mock-derived EVs and conversely the incorporation of 11 proteins (ANXA3, CD59, CLIC1, CLU, DAG1, DARS1, EIF3C, GOLM1, GSTM3, PGAM1, PSMB1), while the rate of exosome secretion remained globally unaltered. This hinted at the modulation of EV composition by the virus. As proof of principle, the analysis of select hits including virion and infected EVs host proteins by dsiRNA led to reduced viral yields in many instances (61% of the tested targets). This suggests that EVs and virions reciprocally modulate each other. Furthermore, this highly positive rate of functional knockdowns validates the relevance of our proteomic pipeline and opens news research avenues to impair HCoV-OC43 replication. However, further work is needed to clarify whether the pool of cellular proteins within the virions and EVs are functionally active and to define their mechanisms of action.

Among the subset of proteins tested, CLU, MFGE8, RAB7A and likely EIF3C seemed the most critical players with over 75% inhibition when these proteins were depleted. Interestingly, CLU (clusterin) is a negative modulator of the membrane attack complement system that HCoV-OC43 harnesses to block innate immunity by recruiting it to the surface of infected cells (80). EIF3C is one of many translational initiation factors that modulate hepatitis C gene expression (81, 82) and conceivably also modulate coronaviruses. MFGE8 is a multifunctional protein that has been associated with tissue repair (83) and binds the SARS-CoV-2 receptors expressed in the placenta (84). RAB7A is a small GTPase that modulates the intracellular vesicular transport of proteins along the endocytic pathway (85) that can be used by some coronaviruses to enter cells (86–88) and is modulated by SARS-CoV-2 (89–92). These proteins thus represent interesting putative targets to modulate coronaviruses.

An elegant proteomics study of SARS-CoV-2 virions recently reported that 120 and 284 cellular proteins were statistically enriched in viruses produced by A549-ACE2 and Calu-3 lung cells respectively (48). Most of those proteins are RNA-binding proteins or modulate RNA metabolism and transport, which is in line with our own findings. The authors further showed that the virion-embedded G3BP1 and G3BP2 stress granule nucleators may promote the formation of virion assembly sites, delineating the importance of those virion-associated proteins. Interestingly, eight SARS-CoV-2 virions-associated hits were also found in our study (AGRN, ANXA11, DARS1, EIF5A, FAM3C, HSPG2, PABPC1, RAB6A), pointing to the putative conservation of these host proteins in coronavirus mature virions. Overall, this study not only provides new and important insight into the incorporation of numerous host proteins in HCoV-OC43 virions but also demonstrates their potential impact on the virus and EV composition. The functional analysis of selected proteins by siRNA led to reduced viral yields in many cases, indicating a reciprocal modulation between EVs and virions. This confirms the biological relevance of our proteomic approach and opens new avenues for research to impede HCoV-OC43 replication. Further investigations are necessary to determine whether the host proteins within virions and EVs are functionally active and to elucidate their mechanisms of action.

## Materials and Methods

### Cell lines

The male human ileocecal adenocarcinoma tumour cell line HRT-18 cell line was used for the HCoV-OC43 infection and virus quantification (gift from Dr. Talbot, INRS (93)). Please note that this cell line is distinct from ATCC CCL-244, which is listed as HCT-8/HRT-18. It was cultured in Dulbecco modified Eagle medium (DMEM) with 10% fetal bovine serum (FBS), 1% L-glutamine (G7513; Sigma-Aldrich) and 1% penicillin/streptomycin (P4333; Sigma-Aldrich). Male Vero-E6 cells (ATCC CL-1586) were propagated in DMEM with 5% bovine growth serum (BGS), 1% L-glutamine, and 1% penicillin/streptomycin. When producing virions, the above media was replaced by serum-free media OptiPRO^TM^ (SFM-OptiPRO; 12309019; Gibco).

### Virus

The HCoV-OC43 variant VR-1558 (ATCC, Manassas, VA, USA) was used throughout this study. HRT-18 cells grown in three 600 cm^2^ Corning™ dishes until 80% confluent were mock-treated or infected with the HCoV-OC43 virus at a multiplicity of infection (MOI) of 0.1. After a 1h adsorption at 37°C and 5% CO_2_, 70 ml of SFM-OptiPRO media was added to each dish and incubated at 33°C, 5% CO_2_ for 3 days. Viruses released into the supernatant were harvested and concentrated by one of two means (see below).

### Virus concentration

#### A) Amicon filter

The supernatant of infected cells was collected 72 h post-infection and centrifuged at 800 *x g* for 5 minutes followed by filtration with a 0,45 µm filter to remove debris. The supernatant from the three large dishes were combined and the extracellular virions (and EVs) concentrated on Amicon® Ultra-15 centrifugal filter units with a 100 kDa cut-off (UFC9100; Millipore). The residual concentrated material was topped up to 3 ml with 1x PBS (13.7 mM NaCl, 0.27 mM KCl, 0.2 mM KH_2_PO_4_, 1 mM Na_2_HPO_4_) and stored at −80°C.

#### B) Ultracentrifugation

As a second option, the above cleared supernatant was ultracentrifuged at 20 000, 60 000 or 100 000 *x g* / 4°C for 60 min in thin walled polyclear Seton tubes (Seton scientific; NC9863486) in a P40ST rotor and Hitachi ultracentrifuge (CP-100NX). The pellets were gently resuspended in 200 µl of 1x PBS overnight at 4°C then stored at −80°C.

#### Virus and exosome separation

The separation protocol using iodixanol/sucrose was essentially that of Dogammatzis et al., who successfully reported the separation of HSV-1 viral particles from EVs (70). Briefly, 8-25% iodixanol continuous gradients (Optiprep; Sigma; D1556) in 10 mM Tris (pH 8) and 0.25 M sucrose were prepared with a Biocomp gradient station. One ml of concentrated virions/EVs was loaded on top of each gradient and ultracentrifuged in a Hitachi (CP-100NX) using a P40ST rotor at 250,000 g for 135 min. Eighteen 600 µl fractions were manually collected from the top to the bottom of the gradient using a peristaltic pump.

#### Western blot and antibodies

Equal volumes (50 µl) of each gradient fraction were mixed with loading buffer (final concentration: 50 mM Tris-HCl, pH 6.8, 2% SDS, 0.1% bromophenol blue, 10% glycerol, and 2% β-mercaptoethanol) and heated at 95°C for 5 minutes. Samples were then loaded and separated on 5 to 20% SDS-PAGE gradient gels, then transferred to PVDF membranes (Bio-rad) and probed with specific primary antibodies (see below) along with HRP-linked secondary antibodies. Finally, the membranes were revealed using ECL (170-5060; Bio-rad) and a Chemidoc (Bio-Rad). Primary antibodies for Western blotting (WB) were from the following sources and dilutions: Rabbit monoclonal anti-CD9 (1:1000; Cell Signalling, #13174), rabbit monoclonal anti-CD63 (1:1000, Abcam, 134045), mouse monoclonal antibody anti-CD81 (1:500, Santa Cruz; SC-23962) and mouse monoclonal anti-HCoV-OC43 nucleocapsid protein (1:1000, Millipore Sigma, MAB9012). Secondary antibodies coupled to HRP were purchased from molecular probes and Jackson Immuno-research.

#### HCoV-OC43 quantification using the TCID50-IPA method

The TCID50-IPA assay (median tissue culture infectious dose by immunoperoxidase staining) is a sensitive and reliable method for HCoV-OC43 titration (66). Briefly, three days before infection, HRT-18 cells were seeded in 96-well plates at 7,000 cells/well in DMEM with 10 % FBS at 37°C with 5% CO_2_. On the infection day, the media was removed, cells were infected with 50 µl of a 10-fold serial dilution of the virus with 3 technical replicates and incubated at 33°C, 5% CO_2_. Three days post-infection (DPI) the media was discarded, cells were washed with 1x PBS and fixed with 100% methanol containing 0.3% (v/v) hydrogen peroxide for 15-30 minutes at room temperature followed by incubation with the 4.3E4 mouse monoclonal antibody against the HCoV-OC43 spike protein (dilution 1:50, gift from Dr. Talbot, INRS) and incubation with secondary HRP-goat anti-mouse IgG (H+L) (Jackson Laboratory, Bar Harbor, ME, USA) (1:2000 dilution). Finally, plates were washed 3 times with 1x PBS and incubated for 15 min at RT with 0.03-0.04% (w/v) 3.3′Diaminobenzidine (DAB) containing 0.01% (v/v) H_2_O_2_. The wells were scored as either positive or negative using an Evos XL Core microscope with a 20X objective (Invitrogen, Waltham, MA, USA) and titers calculated according to Spearman and Karber (94, 95).

#### Silver staining

Equal volumes (50 µl) of gradient fractions were loaded onto 5 to 20% SDS-PAGE gradient gels. Following electrophoresis, the gels were incubated for 1 h in a fixative buffer (10% acetic acid; 40% methanol) and washed in demineralized water (dH_2_O). Next, the gels were activated by adding staining activator (0.02% sodium thiosulphate in dH_2_O) for 1-2 minutes. Gels were then rinsed with dH_2_O. The silver solution (0.1% (w/v) silver nitrate, 0.04% (v/v) formaldehyde in dH_2_O) was next added to the gels for 20 min on a shaker. The gels were quickly rinsed with dH_2_O and developed in 2% (w/v) sodium carbonate, 0.04% (v/v) formaldehyde, and 5% (v/v) staining activator in dH_2_O. Gels were then washed with dH_2_O, and the stop solution (5% acetic acid) was added for 2 minutes. Finally, gels were imaged using a G:BOX Chemi (SynGene; XQ).

#### Electron microscopy

Purified HCoV-OC43 fractions 6 and 17 were inactivated using 0.8 % paraformaldehyde 1M HEPES pH 7.6 for 30 minutes at 4°C. The inactivated samples were deposited on square 200-mesh Formvar carbonated copper-coated grids (Electron Microscopy Sciences; FCF200-Cu-50) and contrasted using 2% uranyl acetate (Canemco and Maivac). The grids were then washed in 1M HEPES buffer (Sigma; 83264; pH: 7.6) and dried. Samples were examined with a Philips Tecnai 12 Transmission Electron Microscope.

#### Mass spectrometry

The protein concentration of gradient fractions 6 and 17 was quantified using the Pierce BCA Protein Assay Kit (Thermo Fisher). Ten micrograms of the purified fractions 6 (exosome fraction) and 17 (virus fraction) from both mock and HCoV-OC43-infected samples were inactivated by heating 1h at 80°C. The samples were reconstituted in 50 mM ammonium bicarbonate with 10 mM TCEP (Tris(2-carboxyethyl)phosphine hydrochloride; Thermo Fisher Scientific), and vortexed for 1 h at 37°C. For alkylation, chloroacetamide (Sigma-Aldrich; final concentration of 55 mM) was added. Samples were vortexed for another hour at 37°C. Protein digestion was conducted by addition of 1 µg of trypsin for 8 h at 37°C. Samples were dried down and solubilized in 5% acetonitrile (ACN) −4% formic acid (FA). Samples were loaded on a 1.5 µl pre-column (Optimize Technologies, Oregon City, OR). Peptides were separated on a reversed-phase column (150-μm i.d. by 200 mm) with a 56 min gradient from 10 to 30% ACN-0.2% FA and a 600 nl/min flow rate on an Easy nLC-1200 connected to an Exploris 480 (Thermo Fisher Scientific, San Jose, CA). Each full MS spectrum acquired at a resolution of 120,000 was followed by tandem-MS (MS-MS) spectrum acquisition on the most abundant multiply charged precursor ions for 3s. Tandem-MS experiments were performed using higher energy collision dissociation (HCD) at a collision energy of 34%. The data were processed using PEAKS X Pro (Bioinformatics Solutions, Waterloo, ON) and a concatenated database made of Uniprot human (20349 entries) and HCoV-OC43 (22 entries) databases. Mass tolerances on precursor and fragment ions were 10 ppm and 0.01 Da, respectively. Fixed modification was carbamidomethyl (C) while variable modifications were acetylation (N-ter), oxidation (M), deamidation (NQ), phosphorylation (STY). The data were visualized with Scaffold 5.1.2 (95% protein and peptide thresholds; minimum of 2 peptides identified).

#### DsiRNA knock-down

A total of 170,000 HRT-18 cells per well were seeded in 12-well plates 24 h before transfection. One hour before transfection, the media was changed to SFM-OptiPRO. Cells were transfected with the LipoJet siRNA transfection kit (SignaGen #SL100468) and 100 nM dsiRNA pools of two distinct dsiRNAs per target, except for PRSS2, for which a single dsiRNA was proposed by IDT (Table 3). Twenty-four hours post-transfection, HRT-18 cells were infected with HCoV-OC43 at a MOI of 0.1. Forty-eight hours post-infection in OptiPro media (72 h post-transfection), cells were collected in 200 µl PBS and centrifuged for 10 minutes at 315 *x g* at 4°C. They were next sonicated on ice 15 times for 1 second at intensity 8 in a Sonic Dismembrator Model 100 sonicator equipped with a cup horn. After four cycles of freeze-thawing, they were stored at −80°C. Virus titration was performed by the TCID50-IPA method. A non-targeting control, NC1 (IDT DNA; 51-01-14-04), was used.

**Table 3:**
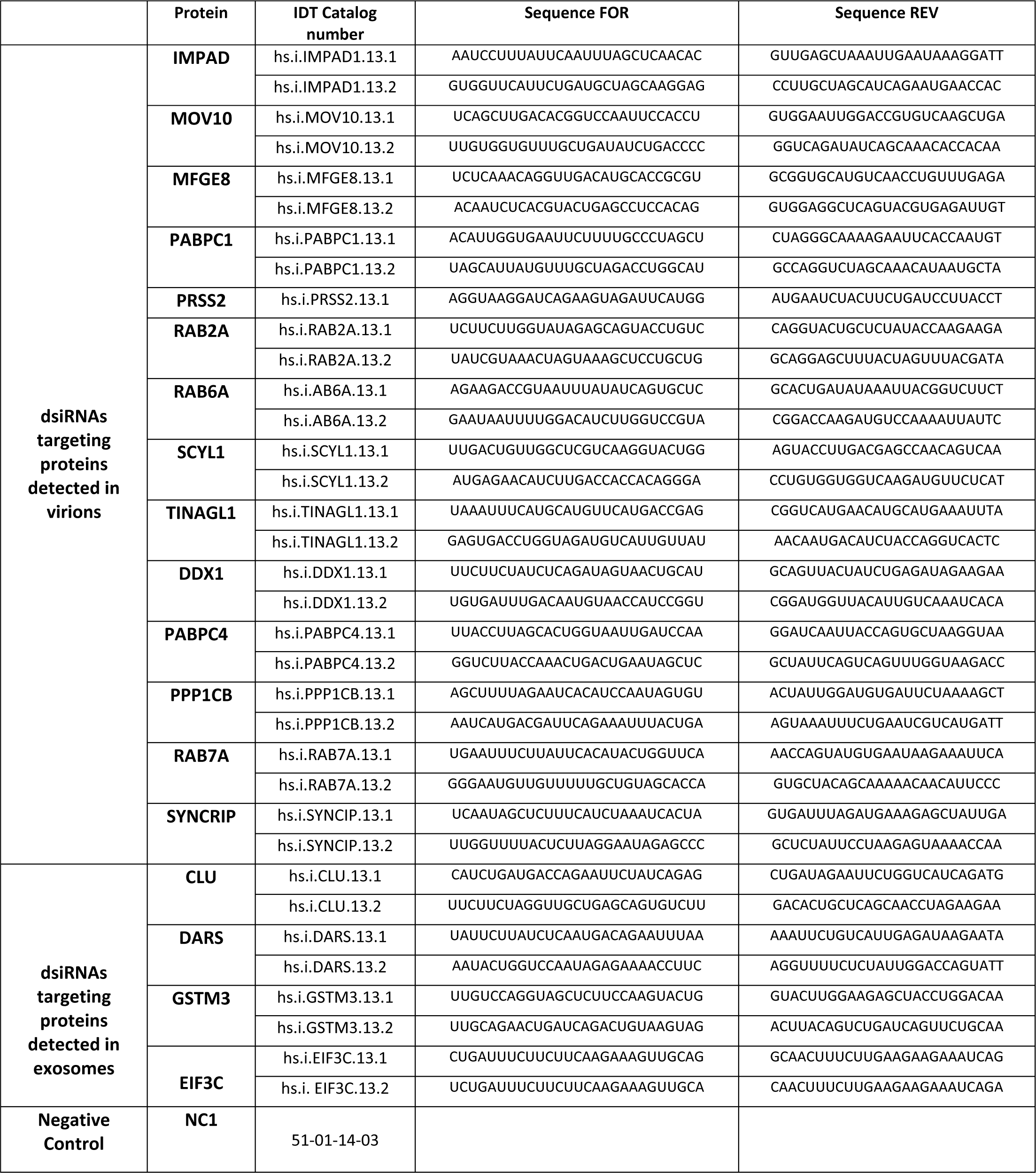
dsiRNA reagents used in this study. With the exception of PRSS2, commercial pools included two distinct dsiRNA per target, as proposed by our supplier (IDT). The NC2 sequence is proprietary and unknown.

#### Viability assay

Twenty-four hours before the viability test, HRT-18 cells were seeded into 96-well Greiner black plates. An hour before transfection, fresh SFM-OptiPRO media was added to all wells. LipoJet siRNA transfection kit (SignaGen #SL100468) was used to transfect the cells with 100 nM of control dsiRNA (NC1) or the dsiRNAs targeting host proteins (see below). Cellular activity was monitored at 72 h post-transfection by adding 10% alamarBlue (Invitrogen; DAL1025). Fluorescence intensity was measured using a BMG Labtech Clariostar microplate reader at 560 nm and 590 nm. All readouts were normalized to the non-transfected control.

## Acknowledgments

We are most indebted to Dr. Marc Desforges and feu Dr. Pierre Talbot for their expertise and for providing reagents. We also wish to thank Dainelys Guadarrama Bello and Dr. Antonio Nanci for help with electron microscopy and Sandrine Marqueteau for the production and titration of the virus. Finally, thanks to all lab members for their useful critical comments. Pilot funding for this research was from a competitive Faculty of Medecine (University of Montreal) and the Sainte-Justine University Hospital Foundation award to RL and NG. The funders had no role in study design, data collection and interpretation or the decision to submit the work for publication.

**Supplementary Tables (Proteins lists):** The different tabs of this Excel file show the cellular proteins associated with mock or HCoV-OC43 infected virions or exosome fraction based after hand curation. The high confidence proteins that were enriched compared to the CRAPome database are shown in bold.

